# PI(4,5)P_2_ forms dynamic cortical structures and directs actin distribution and cell polarity in C. elegans embryos

**DOI:** 10.1101/215079

**Authors:** Melina J. Scholze, Kévin S. Barbieux, Alessandro De Simone, Mathilde Boumasmoud, Camille C. N. Süess, Ruijia Wang, Pierre Gönczy

**Affiliations:** Swiss Institute for Experimental Cancer Research (ISREC), School of Life Sciences, Swiss Federal Institute of Technology (EPFL), Lausanne, Switzerland; Geodetic Engineering Laboratory (TOPO), Swiss Federal Institute of Technology (EPFL), Environmental Engineering Institute (IIE), Lausanne Switzerland

**Keywords:** C. elegans embryo, Phosphoinositiodes, PIP_2_, asymmetric cell division, actin, PAR polarity

## Abstract

Asymmetric division is crucial for embryonic development and stem cell lineages. In the one-cell *C. elegans* embryo, a contractile cortical actomyosin network contributes to anterior-posterior (A-P) polarity and asymmetric division by segregating PAR proteins to discrete cortical domains. Here, we discovered that the plasma membrane lipid phosphatidylinositol 4,5-bisphosphate (PIP_2_) forms dynamic structures in *C. elegans* zygotes, distributing in a polarized and PAR-dependent manner along the A-P axis. PIP_2_ cortical structures overlap with F-actin and coincide with the actin regulators RHO-1, CDC-42 and ECT-2. Particle image velocimetry analysis revealed that PIP_2_ and F-actin cortical movements are coupled, with PIP_2_ structures moving slightly ahead. Importantly, we established that PIP_2_ cortical structures form in an actin-dependent manner and, conversely, that decreasing or increasing the level of PIP_2_ results in severe F-actin disorganization, revealing the interdependence between these components. Furthermore, we uncovered that PIP_2_ regulates the sizing of PAR cortical domains. Overall, our work establishes for the first time that a lipid membrane component, PIP_2_, is a critical modulator of actin organization and cell polarity in *C. elegans* embryos.

**Summary statement:** PI(4,5)P_2_ is distributed in dynamic cortical structures and regulates asymmetric division by controlling actin organization and cell polarity in the one-cell *C. elegans* embryo.

## Introduction

Asymmetric division generates cellular diversity and is particularly prevalent during development. During intrinsic asymmetric division, polarity is established and maintained in a mother cell; thereafter, polarity is translated into correct spindle positioning during mitosis along this polarity axis, resulting in the proper cleavage and partition of cellular contents to daughter cells. The extensively studied and evolutionarily conserved partitioning defective (PAR) proteins are critical for cell polarity and asymmetric division (reviewed in Goldstein and Macara, 2007; Gönczy, 2008; Knoblich, 2010). By contrast to the wealth of knowledge regarding PAR proteins and interacting components, the involvement of lipid plasma membrane components in cell polarity is less understood, in particular in developing systems.

The early *C. elegans* embryo has proven instrumental for dissecting the mechanisms governing asymmetric division (reviewed in Hoege and Hyman, 2013; Pacquelet, 2017; Rose and Gönczy, 2014). Shortly after fertilization, the entire embryo surface exhibits uniform contractions of the cortical actomyosin network located underneath the plasma membrane (Munro et al., 2004). These contractions are driven by non-muscle myosin 2 (NMY-2), which is activated by the Rho GTPase RHO-1 and its guanine nucleotide exchange factor (GEF) ECT-2 (Motegi and Sugimoto, 2006; Schonegg and Hyman, 2006). Sperm-derived centrioles are key for breaking symmetry of this system and for inducing local disappearance of cortical ECT-2 in their vicinity, thereby determining the embryo posterior. This leads to local inactivation of RHO-1 and initiation of cortical flows away from this region, towards the future embryo anterior (Bienkowska and Cowan, 2012; Cowan and Hyman, 2004; Motegi and Sugimoto, 2006). Polarized contractility promotes establishment of PAR polarity, whereby PAR-3, PAR-6, and atypical protein kinase C-like 3 (PCK-3) are segregated to the anterior side, whereas PAR-1, PAR-2, and Larval Giant Larvae-like 1(LGL-1) occupy the expanding posterior cortical domain (reviewed in Hoege and Hyman, 2013; Pacquelet, 2017; Rose and Gönczy, 2014). The RhoGTPase CDC-42 is also segregated to the anterior, where it stabilizes the actomyosin network and promotes PAR-6 association with the cortex (Kumfer et al., 2010; Motegi and Sugimoto, 2006; Schonegg and Hyman, 2006).

PAR polarity can be established in *C. elegans* zygotes also though a partially redundant pathway, whereby microtubules nucleated from centrosomes protect PAR-2 from PKC-3-mediated phosphorylation, thus allowing PAR-2 association with phospholipids at the embryo posterior (Motegi et al., 2011). In other systems, homologues of PAR proteins and interacting components also associate with phospholipids, as exemplified by *Drosophila* DmPar3 binding to phosphatidylinositol 4,5-bisphosphate (PI(4,5)P_2_, referred to hereafter as PIP_2_for simplicity) and phosphatidylinositol 3, 4,5-triphosphate (PI(4,5,6)P_3_, hereafter PIP_3_) (Krahn et al., 2010). Furthermore, human Cdc42 binds to PIP_2_ (Johnson et al., 2012). Overall, whereas it is clear that phospholipids can bind PARs and interacting proteins in some contexts, their subcellular distribution and potential function in an asymmetrically dividing system such as the *C. elegans* zygote remain unclear.

PIP_2_ is the most abundant of seven phosphorylated phosphatidylinositols and is present mostly in the inner leaflet of the plasma membrane, as revealed for instance by the distribution of the pleckstrin homology (PH) domain of mammalian phospholipase C1δ1 (PLC1 δ1), which binds PIP_2_ *in vitro* and in cells with high specificity (Garcia et al., 1995; Lemmon et al., 1995; Várnai and Balla, 1998). PIP_2_ is mainly phosphorylated from phosphatidylinositol 4-phosphate PI(4)P (PIP) by Type I PI(4)P5-kinases (PIP5K1), and can be further phosphorylated to PIP_3_ by Phosphatidylinositol 3-kinases (PI3K). Conversely, PIP_2_ can be dephosphorylated by 5-phosphatases, including OCRL and synaptojanin (reviewed in Brown, 2015; De Craene et al., 2017; McLaughlin et al., 2002). Amongst other roles, in systems from S. *cerevisiae* to *H. sapiens,* PIP_2_ helps link the F-actin cortical network to the plasma membrane, as well as stimulate F-actin assembly and reorganization. The latter function is achieved notably by activating, together with Cdc42, WASP family proteins that, in turn, activate the actin nucleator Arp2/3. Moreover, PIP_3_ further activates WASP family proteins through RhoGTPase GEFs (reviewed in Brown, 2015; De Craene et al., 2017; Di Paolo and De Camilli, 2006; McLaughlin et al., 2002; Wu et al., 2014; Yin and Janmey, 2003; Zhang et al., 2012). Whether and, if so, how, PIP_2_ regulates cortical actomyosin network organization in the *C. elegans* embryo is not known.

Not only is the potential function of PIP_2_ in the *C. elegans* zygote not clear, but the same holds for PIP_2_ subcellular distribution. In other systems, PIP_2_ can distribute unevenly in the plasma membrane, for instance accumulating in macrodomains in nascent phagosomes, in membrane ruffles or at the leading edge of motile cells, which all exhibit curved membranes that are sites of actin reorganization (Chierico et al., 2015; McLaughlin et al., 2002; Zhang et al., 2012). Accordingly, PIP_2_ can stimulate actin polymerization in curved but not flat model membranes (Gallop et al., 2013). Interestingly, PIP_2_ patches assemble at the leading edge of neuronal PC12 cells prior to F-actin patch accumulation, but their formation also depends on F-actin (Golub and Caroni, 2005; Golub and Pico, 2005). Moreover, it was suggested that F-actin enrichment in cell cortices drives clustering of PIP_2_-containing macrodomains, which in turn further regulate actin polymerization and branching (reviewed in Chichili and Rodgers, 2009). Overall, PIP_2_and F-actin polymerization function in a positive feedback mechanism in several systems.

The single PIP5K1 in *C. elegans* is PPK-1, which can synthesize PIP_2_ from PIP *in vitro* and *in vivo* (Weinkove et al., 2008). Overexpression of PPK-1 in developing worm neurons increases the level of PIP_2_ and results in extended filopodial-like structures, probably through changes in the actin cytoskeleton (Weinkove et al., 2008). In the somatic gonad, PPK-1 is important for F-actin cytoskeletal reorganization and, therefore, gonad contractility (Xu et al., 2007). Moreover, PPK-1 is enriched on the posterior cortex of one-cell embryos and has been reported to be important for asymmetric spindle positioning, but not for cell polarity (Panbianco et al., 2008). In other systems, however, PIP_2_ can regulate cell polarity through its ability to recruit PAR proteins and reorganize the actin cytoskeleton. Thus, in the *Drosophila* follicular epithelium, PIP_2_ recruits DmPar3 to the apical plasma membrane to maintain apical-basal polarity (Claret et al., 2014). Moreover, PIP_2_ might mediate interactions between PAR proteins, the actomyosin network and the plasma membrane in the fly oocyte (Gervais et al., 2008), as well as regulate apical constriction in the fly embryo (Guglielmi et al., 2015). Motile cells such as mammalian neurophils or *Dictyostelium discoideum* also rely on PIP_2_, together with PIP_3_, for actin network reorganization and polarization (reviewed in Wu et al., 2014). To summarize, in many systems, PIP_2_ is essential for F-actin reorganization and cell polarization, but it remains to be investigated whether this is the case in the developing *C. elegans* embryo.

## Results

### The PIP_2_ biomarker GFP::PH^PLC1δ1^ is present in dynamic polarized cortical structures in one-cell C. elegans embryos

While monitoring the distribution of components involved in asymmetric division of the *C. elegans* zygote with confocal spinning disk microscopy, we found that the PIP_2_ biomarker GFP::PH^PLC1δ1^ (Audhya et al., 2005)forms distinct and dynamic structures at the cell cortex (Fig. 1A-J; Fig. S1A; Movie 1). Initially, when the cortical actomyosin network is contractile throughout the embryo, PIP_2_ is present weakly and evenly on the cell cortex (data not shown). Thereafter, when the actomyosin network begins to retract towards the anterior at the onset of polarity establishment, we observed the appearance of striking elongated cortical structures enriched in PIP_2_, primarily on the anterior side of the embryo (Fig. 1A, B, 1K; Fig. S1A, top). Such PIP_2_ cortical structures have an average size of ~2.5 µm^2^ and further elongate over time as the cell progresses through the cell cycle (Fig. 1L, M). Subsequently, all elongated PIP_2_ cortical structures move anteriorly so that they become distributed in a clearly polarized manner at pseudocleavage, which marks the end of the polarity establishment phase (Fig. 1C, D, arrow). At this time, PIP_2_ structures cover ç15% of the anterior cortical surface (Fig. 1K). During the centration/rotation stage that follows, PIP_2_ cortical structures first decrease in size (Fig. 1E, F, arrowhead), with most of them disappearing completely but some remaining as small foci by the time the cell enters mitosis (Fig. 1G, H, arrowheads). A few elongated cortical structures reappear during cytokinesis, primarily in the embryo anterior (Fig. 1I, J). In contrast to the discrete structures visible when imaging the cortical plane, PIP_2_ entities are barely detectable in the embryo middle plane (Fig. S1A, bottom), likely explaining why they were not reported previously (Audhya et al., 2005; Blanchoud et al., 2010; Panbianco et al.,2008).

**Fig. 1.**
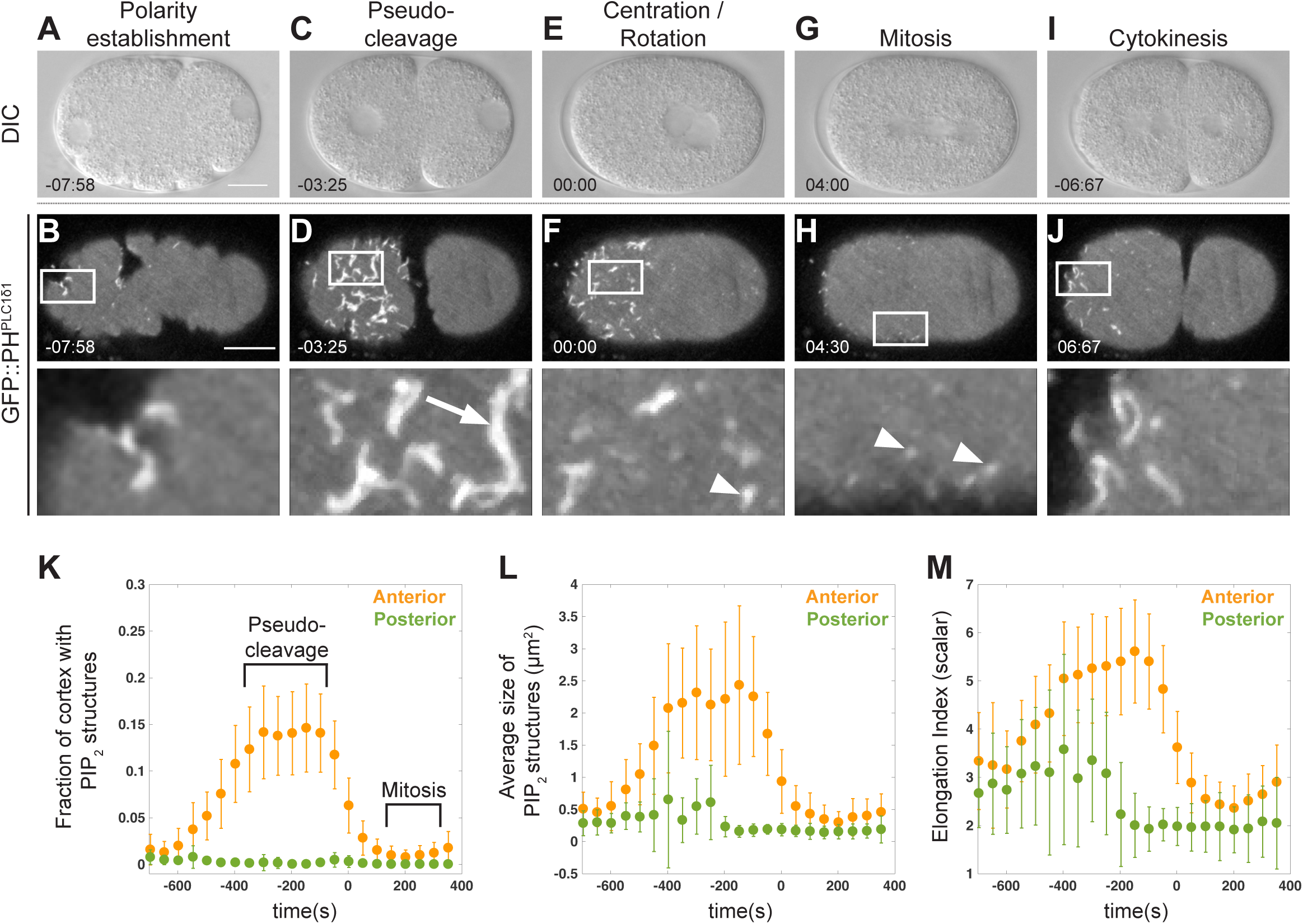
The PIP_2_ biomarker GFP::PH^PLC1δ1^ is enriched in dynamic cortical structures. **(A-J)** Differential interference contrast (DIC) (A, C, E, G, I, middle plane of the embryo) and spinning disk confocal imaging (B, D, F, H, J, cortical plane of a different embryo at the corresponding stages, with boxed regions magnified below) of one-cell *C. elegans* embryos at the indicated stages expressing GFP::PH^PLC1δ1^ monitoring PIP_2_. Unless indicated otherwise, scale bar in this and subsequent figures: 10 µm. Time is indicated in minutes:seconds, with 00:00 corresponding to the time of centration/rotation (to in K-M). See also Movie 1. Here and in all subsequent figures, the embryo anterior is to the left. **(K)** Fraction of cell cortex covered by PIP_2_ structures. The timing of pseudocleavage and mitosis are indicated. Here and in similar subsequent panels: anterior (orange) and posterior (green) quantitative data is shown, with the mean and the standard deviation. N=39 embryos for K-M. **(L)** Average cortical area covered by PIP_2_ structures overtime. **(M)** Elongation Index of PIP_2_ cortical structures over time. Larger values correspond to most elongated shapes.

We set out to verify the cortical distribution revealed by GFP::PH^PLC1δ1^ using fluorescently labeled synthetic PIP_2_. To this end, we delivered Bodipy-FL-PIP_2_ lipids to embryos whose eggshell had been permeabilized using *perm-1(RNAi)* (Carvalho et al., 2011). Although there was a high background of fluorescent lipids outside the embryo, we found that Bodipy-FL-PIP_2_ distributes in cortical structures marked by mCherry::PH^PLC1δ1^ (Fig. S1B, C). Overall, we conclude that PIP_2_ forms dynamic and polarized structures at the plasma membrane of one-cell *C. elegans* embryos.

### A-P polarity cues regulate the polarized distribution of PIP_2_ cortical structures

We set out to address what regulates the polarized distribution of PIP_2_ cortical structures, which is particularly apparent during the pseudocleavage stage, as evidenced also by the fact that they do not overlap with GFP::PAR-2, which marks the posterior cortical domain (Fig. 2A). By contrast, we found that PIP_2_ cortical structures overlap with the anterior polarity domain harboring GFP::PAR-6 (Fig. 2B; Movie 2). Moreover, we observed that PIP_2_ cortical structures overlap only with elongated GFP::PAR-6 cortical structures (Fig. 2B, arrow), but not with GFP::PAR-6 foci (Fig. 2B, arrowhead), which are two distinct cortical populations of GFP::PAR-6 previously reported to exist (Beers and Kemphues, 2006; Robin et al., 2014; Rodriguez et al., 2017; Wang et al., 2017).

**Fig. 2.**
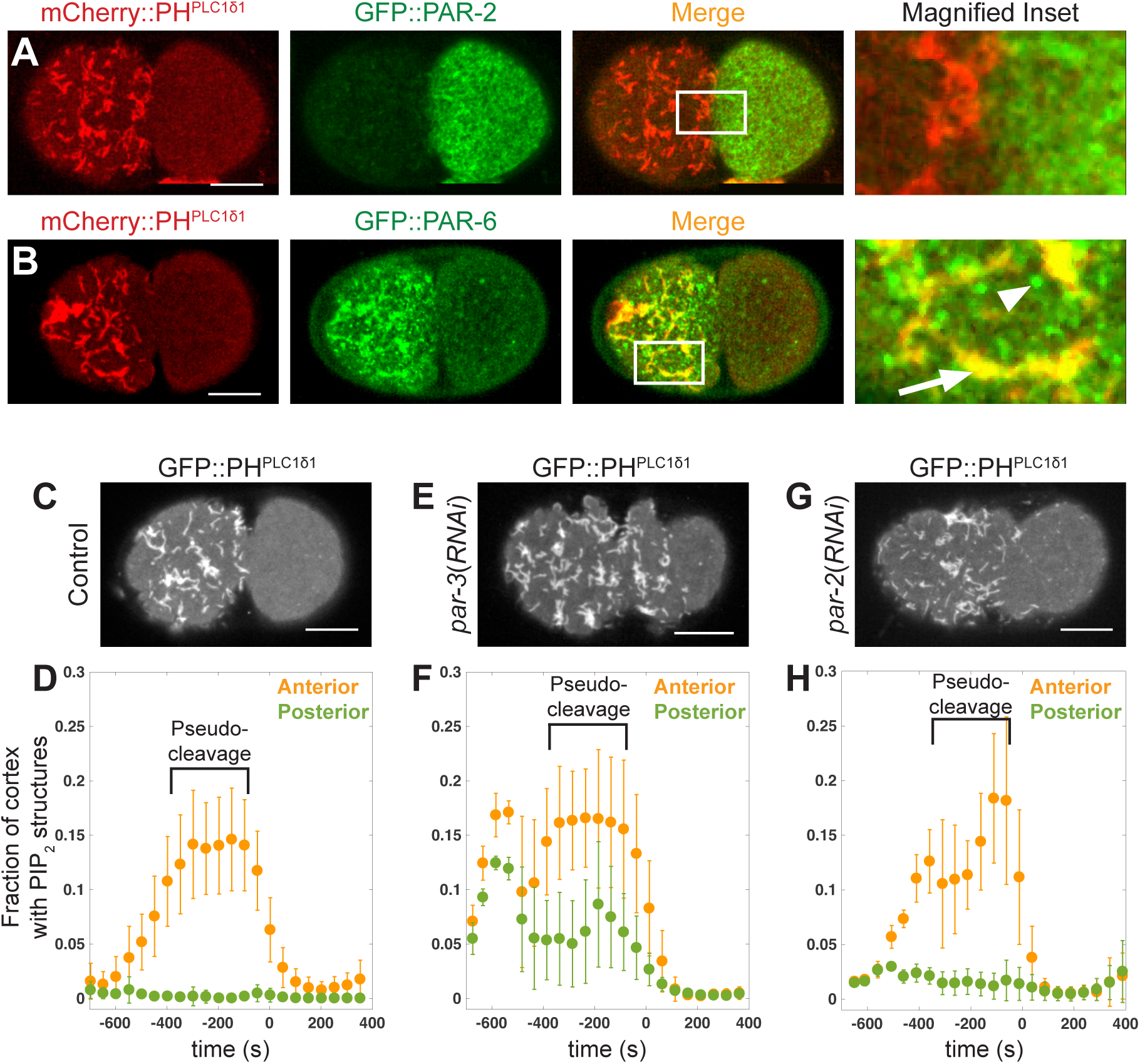
PIP_2_ cortical structures depend on A-P polarity. **(A, B)** Dual color spinning disk confocal cortical imaging of *C. elegans* embryos at the pseudocleavage stage harboring the indicated pairs of fusion proteins; the column on the right shows high magnification views of the boxed regions. (A) mCherry::PH^PLC1δ1^ and GFP::PAR-2, N=5. (B) mCherry::PH^PLC1δ1^ and GFP::PAR-6, N=5; note that elongated cortical structures (arrowhead) but not foci (arrow) of GFP::PAR-6 overlap with mCherry::PH^PLC1δ1^ cortical structures. See also Movie 2. **(C, E, G)** Cortical plane images at pseudocleavage from movies acquired by spinning disk confocal imaging of control (C), *par-3(RNAi)* (E, N=10) or *par-2(RNAi)* (G, N=11) *C. elegans* embryos expressing GFP::PH^PLC1δ1^. **(D, F, H)** Fraction of cell cortex covered by PIP_2_ structures on the anterior (orange) and posterior (green) side in the conditions corresponding to (C, E, G).

We next tested whether the polarized distribution of PIP_2_ cortical structures depends on A-P polarity cues. In contrast to the polarized distribution observed in the control condition, we found that upon *par-3(RNAi),* PIP_2_ cortical structures distribute essentially uniformly over the cell cortex (compare Fig. 2C, D with 2E, F), except for the very posterior of the embryo, consistent with the known slight posterior clearing of the actomyosin network upon PAR-3 inactivation (Kirby et al., 1990; Munro et al., 2004). Furthermore, we found that upon *par-2(RNAi),* PIP_2_ cortical structures first move anteriorly (Fig. 2G, H), but then become distributed in a more uniform manner (Fig. 2H), in line with PAR-2 being dispensable for polarity establishment, but essential for polarity maintenance (Cuenca et al., 2003; Hao et al., 2006; Munro et al., 2004). Together, these findings establish that the asymmetric distribution of PIP_2_ cortical structures is regulated by PAR-dependent A-P polarity cues.

### PIP_2_ cortical structures colocalize partially with actin and fully with ECT-2, RHO-1 and CDC-42

Since PIP_2_ and F-actin are interdependent in many systems, we tested whether these two components overlap at the cell cortex of *C. elegans* embryos. As shown in Figure 3A and Movie 3, we found a partial overlap of Lifeact::mKate-2, which monitors F-actin, and of PIP_2_ cortical structures marked by mNeonGreen::PH^PLC1δ1^ (mNG::PH^PLC1δ1^). By contrast, we detected no substantial overlap between mCherry::PH^PLC1δ1^ and GFP::NMY-2 (Fig. 3B; Movie 4), the non-muscle myosin that powers contractions of the cortical actomyosin network (Guo and Kemphues, 1996; Munro et al., 2004). Strikingly, in addition, we found that PIP_2_ cortical structures marked by mCherry::PH^PLC1δ1^ fully colocalize with GFP::ECT-2, GFP::RHO-1 and GFP::CDC-42 (Fig. 2C-E; Movie 5). Overall, we conclude that PIP_2_ cortical structures colocalize partially with F-actin, as well as completely with the actomyosin network regulators ECT-2, RHO-1 and CDC-42 in the one-cell *C. elegans* embryo.

**Fig. 3.**
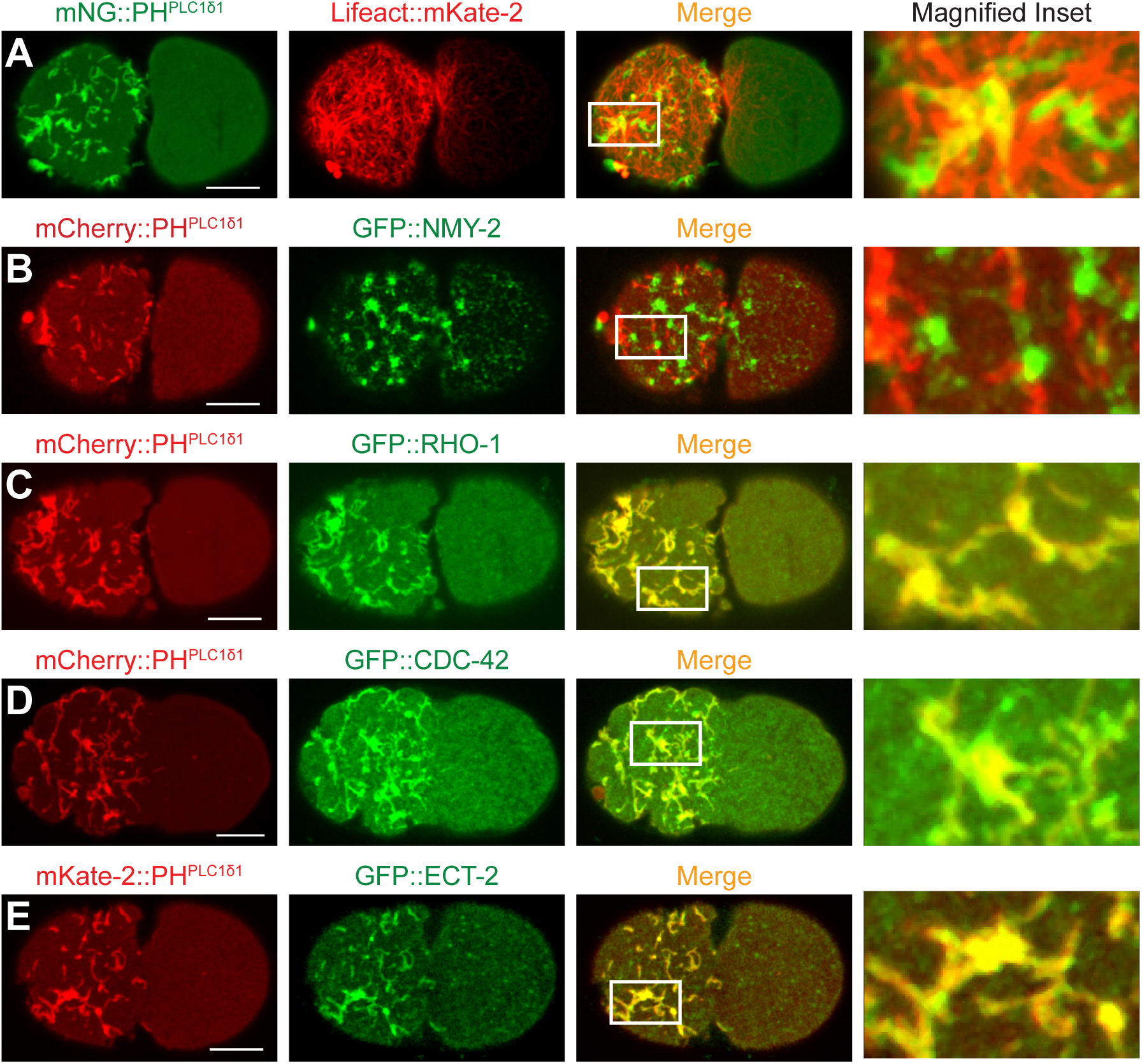
PIP_2_ cortical structures overlap with ECT-2, CDC-42, RHO-1, and partially with actin. **(A-E)** Dual color spinning disk confocal cortical imaging of *C. elegans* embryos at the pseudocleavage stage harboring the indicated pairs of fusion proteins; the column on the right shows high magnification views of the boxed regions. (A) mNeonGreen::PH^PLC1δ1^ and Lifeact::mKate-2, N=46; See also Movie 3. (B) mCherry::PH^PLC1δ1^ and GFP::NMY-2, N=9; See also Movie 4. (C) mCherry::PH^PLC1δ1^ and GFP::RHO-1, N=9; (D) mCherry::PH^PLC1δ1^ and GFP::CDC-42, N=7; See also Movie 5. (E) mKate2-PH^PLC1δ1^ and GFP::ECT-2, N=13.

### PIP_2_ cortical structures and the F-actin cytoskeleton move in concert

Live imaging of embryos expressing both Lifeact::mKate-2 and mNG::PH^PLC1δ1^ suggested that movements of PIP_2_ cortical structures and of the F-actin network are somehow coupled, as drastic changes in PIP_2_ cortical structures coincide with alterations in the actomyosin network across the first cell cycle (Movie 3). To investigate this potential coupling in a quantitative manner, we used particle image velocimetry (PIV) to analyze cortical flows of Lifeact::mKate-2 and mNG::PH^PLC1δ1^ in the same embryo during polarity establishment (Fig. 4A-C) (Thielicke and Stamhuis, 2014). This analysis revealed highly correlated local flow velocities at each time point (Fig. 4B; Fig. S2A; Pearson coefficient ρ = 0.61, p<0.0001). Moreover, the direction of flow vectors at each time point is very similar (Fig. 4C; Fig. S2B; θ_cut off_=38° p<0.0001, Material and Methods). Analogous findings were made when comparing mCherry::PH^PLC1δ1^ and GFP::CDC-42 (Fig. S2C). By contrast, no strong correlation was found between mCherry::PH^PLC1δ1^ and the caveolin marker CAV-1::GFP (Fig. S2C), which has been suggested to mark lipid rafts in *C. elegans* (Entchev and Kurzchalia, 2005; Kurzchalia and Ward, 2003; Kurzchalia et al., 1999; Merris et al., 2003). Together, these results demonstrate that cortical movements of PIP_2_ cortical structures and the F-actin network are coupled.

**Fig. 4.**
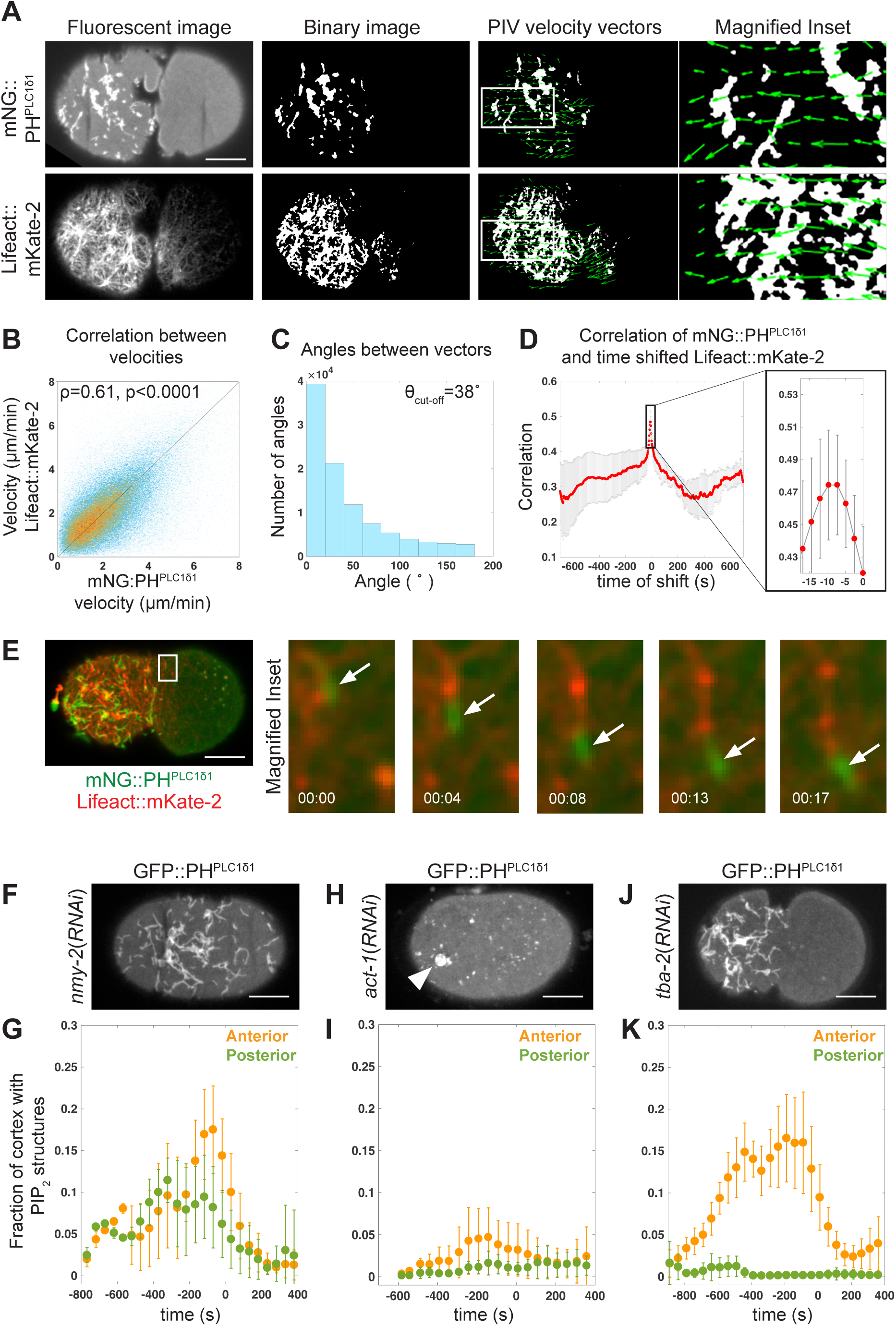
Coupling between PIP_2_ cortical structures and F-actin. **(A)** First column: representative fluorescent images of spinning disk confocal cortical imaging of embryos expressing mNG::PH^PLC1δ1^ and Lifeact::mKate-2 used to perform the PIV analysis. Second column: binary images of thresholded fluorescent images. Third column: PIV velocity vectors (high magnification views of the boxed regions on the right), arrow direction and length represent flow direction and velocity, respectively. **(B)** mNG::PH^PLC1δ1^ flow velocity plotted as a function of Lifeact::mKate-2 velocity in the same position and at the same time. Data points are represented with a color scale dependent on their spatial density, from denser to sparser (red, yellow, light blue). Pearson correlation coefficient: p=0.61, p=0 with Matlab precision, Student’s t test. N=13 embryos for B-D. **(C)** Angle distribution between flow velocity vectors of mNG::PH^PLC1δ1^ and Lifeact::mKate-2 in the same position and at the same time. The angle distribution peaks at 0 = 0° and decays exponentially thereafter (cutoff angle: 0=38°). Two independent velocity fields cannot result in the observed angle distribution (probability: p=0 with Matlab precision (shi2-test)). **(D)** Cross-correlation between thresholded binary movies of mNG::PH^PLC1δ1^ and Lifeact::mKate-2; Lifeact::mKate-2 was shifted with different time intervals relative to mNG::PH^PLC1δ1^. The boxed region is magnified on the right, showing that average maximal overlap is achieved with a time shift of At = −9.3 +/−1.5 seconds (average and standard deviation), irrespective of the order in which the two signals were recorded (Materials and Methods). See also Movie 6. **(E)** Embryo expressing mNG::PH^PLC1δ1^ and Lifeact::mKate-2 imaged every 4.2 seconds; the boxed region is magnified on the right and shows snapshots from corresponding movie illustrating that mNeonGreen::PH^PLC1δ1^ (white arrow) moves ahead of Lifeact::mKate-2. **(F, H, J)** Cortical plane images at pseudocleavage from spinning disk confocal imaging of *C. elegans* embryos treated as indicated and expressing GFP::PH^PLC1δ1^. (F) *nmy-2(RNAi),* N=7; (H) *act-1(RNAi),* N=8 at pseudocleavage and N=12 at mitosis; (J) *tba-2(RNAi),* N=10. Note that both *act-1(RNAi)* and *tba-2(RNAi)* correspond to severe but partial depletion conditions, as more complete depletion results in sterility. Arrowhead: remaining PIP_2_ structure. **(G, I, K)** Fraction of cell cortex covered by PIP_2_ structures on the anterior (orange) and posterior (green) of *nmy-2(RNAi)* (G, N=7), *act-1(RNAi)* (I, N= 8 at pseudocleavage, N=12 at mitosis) or *tba-2(RNAi)* (K, N=10) embryos.

We next addressed whether the movements of PIP_2_ cortical structures and of F-actin are synchronous or instead exhibit a time shift that might suggest a potential hierarchy between them, with one component leading the other. Close examination of movies of embryos expressing Lifeact::mKate-2 and mNG::PH^PLC1δ1^ suggested that PIP_2_ cortical structures move slightly ahead of cortical F-actin (Fig. 4E; Movie 6). To address this possibility quantitatively, we cross-correlated time-shifted images of Lifeact::mKate-2 and of mNG::PH^PLC1δ1^ to determine when there is maximal overlap between the two signals. As shown in Figure 4D, this analysis revealed that maximal correlation is achieved when mNeGr::PH^PLC1δ1^ is on average ~9.3 +/−1.5 seconds ahead of Lifeact::mKate-2. Overall, we conclude that PIP_2_ cortical structures move together with, but slightly ahead of, F-actin filaments.

What drives these movements of PIP_2_ cortical structures? The observation that PIP_2_ cortical structures move slightly ahead of F-actin filaments, with the trailing end of PIP_2_ cortical structure being seemingly in contact with the leading tip of coupled actin filaments (see Fig. 4E), led us to hypothesize that actin polymerization might push PIP_2_ cortical structures. Compatible with this possibility, we found that the average velocity of PIP_2_ cortical structures was ~0.17 +/−0.03 µm/s (Fig. S2D, E), in the range of actin polymerization driven motility in other systems (Brangbour et al., 2011; Cameron et al., 2000; Carlsson, 2003; Carlsson, 2010; Mogilner and Oster, 1996). These findings lead us to propose that the movements of PIP_2_ cortical structures are driven by actin polymerization.

### PIP_2_ cortical structures depend on F-actin

Given notably the tight coupling between cortical PIP_2_ structures and F-actin, we investigated whether components of the actomyosin network regulate the presence of PIP_2_ entities. As shown in Figure 4F, we found that PIP_2_ cortical structures form and move unabated in *nmy-2(RNAi)* embryos, in which actomyosin network contractility is abolished, although they distribute symmetrically, as expected from the known requirement of NMY-2 in A-P polarity (Fig. 4F, G). Therefore, formation and movement of PIP_2_ cortical structures do not depend on a contractile actomyosin cortex. In stark contrast, we found that PIP_2_ cortical structures hardly form in *act-1(RNAi)* embryos (Fig. 4G, H, I; Fig. S3A-D). Moreover, acute impairment of F-actin through treatment of *perm-1(RNAi)* embryos with the actin polymerization inhibitor Cytochalasin D led to the disappearance of PIP_2_ cortical structures (Fig. S3E, F). By contrast, we found that PIP_2_ cortical structures are essentially independent of the microtubule cytoskeleton, impaired here using *tba-2(RNAi)* (Fig. 4J, K). Overall, we conclude that the formation of PIP_2_ cortical structures depends on F-actin.

### Lowering the cellular level of PIP_2_ impacts F-actin distribution

We set out to address whether, conversely, PIP_2_ regulates F-actin organization. If this were the case, then changing the level of PIP_2_ Ishould alter actomyosin network organization. We tested this prediction first by depleting PIP_2_. To this end, we activated phospholipase C, an enzyme that cleaves PIP_2_, by delivering Ionomycin and Ca^2+^ into *perm-1(RNAi)* embryos (Fig. S4A) (Hammond et al., 2012; Várnai and Balla, 1998). Cleavage of PIP_2_ at the plasma membrane was monitored by the gradual loss of GFP::PH^PLC1δ1^ plasma membrane signal, which enabled us to determine, in each embryo, a comparable time t_1/2_ when half of the initial GFP::PH^PLC1δ1^ plasma membrane fluorescence signal disappeared (Fig. S4B, C). A single plane in the middle of embryos expressing GFP::PH^PLC1δ1^ and Lifeact::mKate-2 was monitored in these experiments, as this proved most reliable for determining t_1/2_. In doing so, we found that PIP_2_ removal following addition of Ionomycin/ Ca^2+^ during pseudocleavage led to rapid alteration of embryo shape on the anterior side, coincident with altered F-actin organization (compare Fig. 5A and 5B, Fig. S4E). An analogous alteration in F-actin distribution was observed following Ionomycin/ Ca^2+^ addition during mitosis (Fig. 5C; Movie 7).

**Fig. 5.**
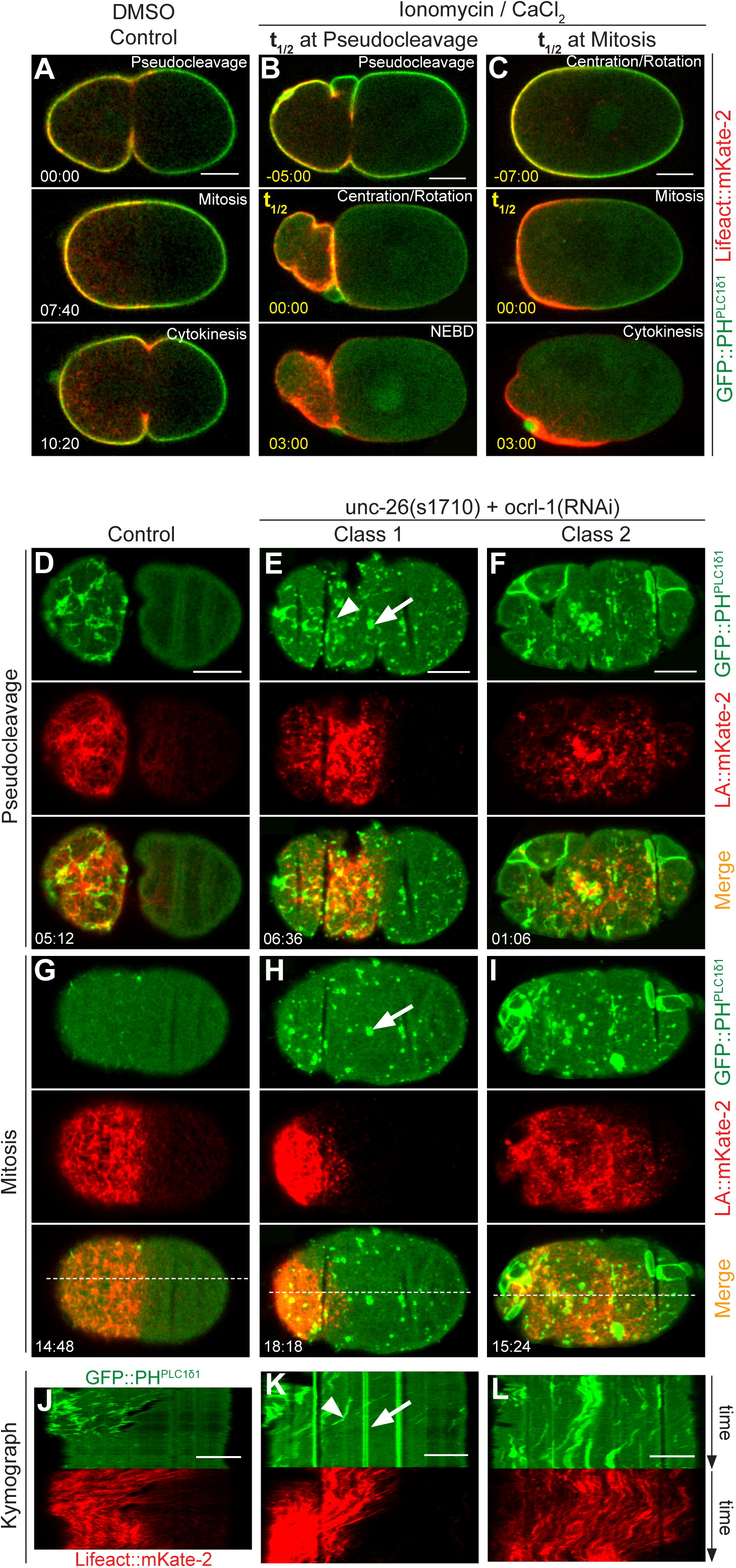
Proper PIP_2_ cellular level is essential for correct organization of the actin cytoskeleton **(A-l)** Confocal spinning disk imaging of embryos expressing GFP::PH^PLC1δ1^ and Lifeact::mKate-2 (A-C: middle plane, D-l: cortical plane). **(A)** DMSO treated *perm-1(RNAi)* control embryos. **(B, C)** *perm-1(RNAi)* embryos treated with Ionomycin/Ca^2+^. t_½_=00:00: time at which half of plasma membrane GFP::PH^PLC1δ1^ fluorescence disappeared; the time stamps are shown in yellow here and in following figure panels where t_½_=00:00. N=17, all stages combined. (B) t_½_ at pseudocleavage. Note that the absence of coverslip, which is needed to preserve fragile *perm-1(RNAi)* embryos, prevents their flattening, such that they are more contractile. Note also that the pseudocleavage furrow moves towards the anterior and either remains there until the end of the first cell cycle (N=4, as shown) or relaxes (N=4, not shown) (C) t_½_ at mitosis. Note that whereas embryos were dissected in the drug-containing solution, drug action (as monitored by t_½_) took place > 6 min after polarity was established. See also Movie 7. **(D-l)** Control (D, G) and *ocrl-1(RNAi) unc-26(s1710)* (E, F, H, I) embryo during pseudocleavage (D, E, F) or mitosis (G, H, I). (E, H) class 1 phenotype (GFP::PH^PLC1δ1^: N=27/48, Lifeact::mKate-2: N=4/11); arrow: immotile structure, arrowhead: motile structure. (F, I) class 2 phenotype (GFP::PH^PLC1δ1^: N=21/48, Lifeact::mKate-2: N=7/11). White dashed line in (G-l) show the position utilized to create the corresponding kymographs in (J-L). See also Movie 9. **(J-L)** Kymographs corresponding to the above movies aligned at cytokinesis arrow: immotile structure, arrowhead: motile structure; Note that motile cortical PIP_2_ structures eventually move towards the cleavage furrow, which partly corrects the aberrant PIP_2_ cortical domain distribution, consistent with the presence of a mechanism correcting mispositioned cortical domains operating at this stage (Schenk et al., 2010). Entire durations of the kymographs, in min:sec: (J) 16:30, (K) 20:00, (L) 19:00.

To test whether the shape change observed following lonomycin/ Ca^2+^ addition is caused by alterations in F-actin organization, we combined this treatment with the actin depoiymerizing agent Latruncuiin A. As shown in Figure S4F and Movie 8, we found that this results in normally shaped embryos. Therefore, shape changes following PIP_2_ removal are F-actin dependent. Together, these results uncover that a normal PIP_2_ level is critical for the proper distribution of F-actin and thus for proper shape of the *C. elegans* zygote.

### Increasing the cellular level of PIP_2_ also impacts F-actin distribution

We sought to test the relationship between PIP_2_ and F-actin further by increasing the level of PIP_2_. We investigated whether this could be achieved by altering individual enzymes from the PIP_2_ biosynthetic pathway using RNAi or mutant worms, but failed to find a single condition where this would be the case (see Table S1 for genes targeted in this study). Therefore, we set out to jointly inactivate OCRL-1 and UNC-26 for the following reasons (Fig. S4A). OCRL-1 is an inositol 5-phosphates that hydrolyzes PIP_2_to phosphatidyl 4-phosphate (PIP), and whose depletion leads to increased level of PIP_2_ on *C. elegans* phagosomes (Cheng et al., 2015). Moreover, UNC-26 is the *C. elegans* homologue of Synaptojanin, a polyphosphoinositide phosphatase that can also hydrolyze PIP_2_ to PIP, and whose impairment results in vesicle trafficking defects and cytoskeletal abnormalities in the worm nervous system (Charest et al., 1990; Harris et al., 2000).

We jointly depleted the function of these two PIP_2_ phosphatases, using RNAi for *ocrl-1* and an extant mutant for *unc-26.* Importantly, we found by comparing cortical mCherry-PH^PLC1δ1^ mean intensity values that *ocrl-1(RNAi) unc-26(s1710)* embryos exhibit an increased overall level of PIP_2_ (Fig. S5A, B). Importantly, in addition, we found that this leads to drastic alterations in PIP_2_ and F-actin cortical structures monitored by GFP::PH^PLC1δ1^ and Lifeact::mKate-2 (compared Fig. 5D and Fig. 5E-F; Fig. S5C-H). First, in addition to motile PIP_2_ structures, we found a population of immotile PIP_2_ clusters residing between the eggshell and the plasma membrane (Fig. 5E, H, K, arrow; Fig. S5I, arrowhead), potentially corresponding to PIP_2_ boluses removed from the cell in an attempt to return to homeostatic conditions. Second, we found that motile PIP_2_ structures do not disappear as readily after pseudocleavage as they do normally (Fig. 5H, I, compare to Fig. 5G; Fig. S5D). Third, we found that motile PIP_2_ structures exhibit altered distribution in all *ocrl-1(RNAi) unc-26(s1710)* embryos (Fig. 5E, F; Fig. S5F, H). In some (hereafter referred as class I embryos, N=27/48), the anterior movements of PIP_2_ cortical structures and of the actomyosin network do not stop at pseudocleavage, but instead continue until the end of mitosis, resulting in a very small anterior domain of PIP_2_ and F-actin cortical structures (compare to Fig. 5G, J to Fig. 5H, K; Movie 9). In class II *ocrl-1(RNAi) unc-26(s1710)* embryos (N=21/48), weak anteriorly directed movement of cortical PIP_2_ and F-actin is initiated, but both components become distributed throughout the cortexby the end of the first cell cycle, except at the very posterior (Fig. 5I, L). Whereas a clear cytokinesis furrow formed in all class 1 embryos, this was the case in only 8/21 class 2 embryos; this subset exhibited more pronounced anteriorly directed movements than the other 13. Overall, we conclude that the extent of PIP_2_ 5-phosphatases depletion is stronger in class 2 than in class 1 embryos, with the severe phenotype in the former perhaps reflecting an impact on multiple cellular processes. Regardless, these findings establish that an increase in the level of PIP_2_ as achieved in class 1 embryos leads to sustained cortical flows towards the anterior side. Moreover, these results further demonstrate that PIP_2_ regulates actin cytoskeletal organization in one-cell *C. elegans* embryos.

### An appropriate level of PIP_2_ is essential for proper PAR polarity establishment and maintenance

It is well known that the actomyosin network is essential for the establishment phase of A-P polarization of the *C. elegans* zygote (Guo and Kemphues, 1996; Hill and Strome, 1988; Hill and Strome, 1990; Munro et al., 2004; Severson and Bowerman, 2003; Shelton et al., 1999). Given that a proper level of PIP_2_ is essential for correct actomyosin network organization, we tested whether it is also important for A-P polarity. To this end, we investigated the impact of excess PIP_2_ on polarity using *ocrl-1(RNAi) unc-26(s1710)* embryos expressing mCherry::PH^PLC1δ1^ and GFP::PAR-2 or GFP::PAR-6, respectively (Fig. 6A-L). We found that the distribution of GFP::PAR-2 and GFP::PAR-6 domains changes in a manner consistent with alterations in motile PIP_2_ structures and the F-actin network, and this from early on (Movie 10). Thus, for GFP::PAR-6, either a small domain formed on the very anterior (Fig. 6B, E; class 1, N=5/9; Movie 10) or else the fusion protein remained present over the entire cortex, except on the very posterior (Fig. 6C,F class 2; N=4/9). As expected, GFP::PAR-2 distributed in a reciprocal manner, either expanding drastically towards the anterior (Fig. 6H, K; class 1, N=12/22; Movie 11) or else remaining restricted to the very posterior (Fig. 6I, L; class 2, N=10/22). Together, these results indicate that a correct level of PIP_2_ is needed for proper F-actin network localization and, presumably as a consequence, appropriate PAR polarity.

**Fig. 6.**
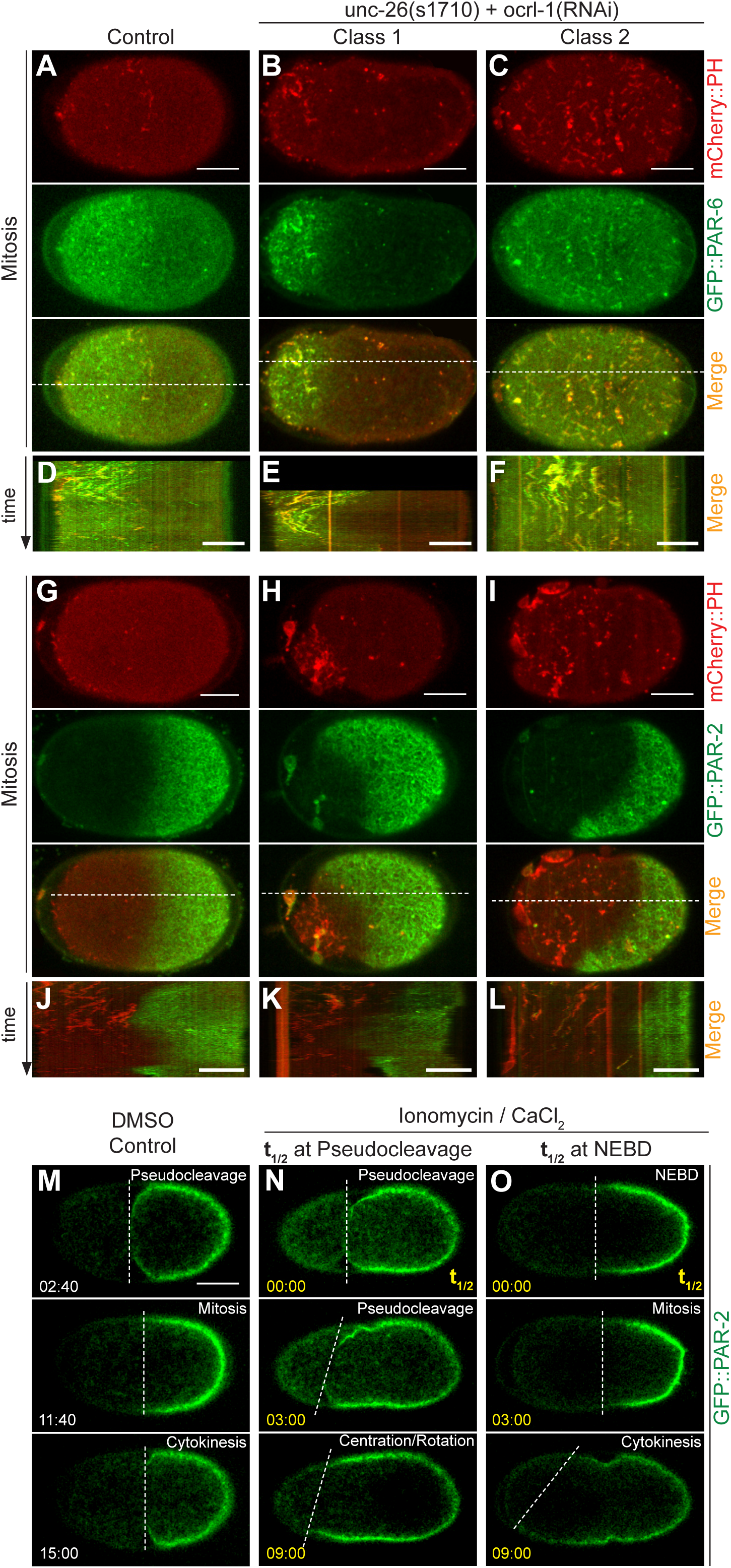
Proper PIP_2_ cellular level is essential for correct PAR polarity. **(A-C, G-l)** Confocai spinning disk cortical imaging of control (A, G) or *ocrl-1(RNAi) unc-26(s1710)* (B, C, H, I) embryos expressing mCherry::PH^PLC1δ1^ and GFP::PAR-6 (A-C, N=5/9 class I, N=4/9 class II) or GFP::PAR-2 (G-I, N=12/22 class I; N=10/22 class II). White dashed lines indicate positions used to create the corresponding kymographs in. See also Movies 10, 11. **(D-F.J-L)** Kymographs corresponding to the above movies aligned at cytokinesis. Entire durations of the kymographs, in min:sec: (D) 16:40, (E) 11:20, (F) 17:40, (G) 17:40, (H) 17:40, (1)17:40. **(M-O)** Images acquired by confocal imaging of embryos expressing GFP::PAR-2, 4.5 µm below the cortical plane. The dashed line marks the boundary of the PAR-2 domain. (M): DMSO treated control *perm-1(RNAi)* embryo. (N, O): *perm-1(RNAi)* embryo treated with Ionomycin/Ca^2+^ (t_½_=00:00) during pseudocleavage, N=6; during mitosis, N=3).Note: worms were dissected in the drug, but drug action (t_½_) took place > 6 min after polarity was established. See Movies 12, 13, 14.

In the above experiments, the level of PIP_2_ was in excess from the beginning of development, such that one could not distinguish whether the impact on polarity reflected a role strictly during the establishment phase or during both establishment and maintenance phases. We reasoned that one could test specifically a potential role for polarity maintenance by adding Ionomycin and Ca^2+^to *perm-1(RNAi)* embryos expressing mCherry::PH^PLC1δ1^ and GFP::PAR-2 after the establishment phase. We found that the GFP::PAR-2 domain is unaltered at t_1/2_ (compare Fig. 6M and 6N, O), even though F-actin is already changed at that moment (see Fig. 5B, C). Importantly, in addition, we found that the GFP::PAR-2 domain expands slowly towards the anterior starting ~3 min thereafter (Fig. 6N, bottom two panels), with the pseudocleavage furrow moving anteriorly initially, and then either retracting (Fig. 6M, N; Movie 12, N=6/14), or else remaining at the very anterior until the end of the first cell cycle Movie 13; N=8/14). Mirroring the findings at pseudocleavage, we found that if t_1/2_ occurs at nuclear envelope breakdown (NEBD), the GFP::PAR-2 domain likewise expands slowly towards the anterior (Fig. 6O; Movie 14). Together these results indicate that an appropriate level of PIP_2_ is essential for proper PAR polarity also during the maintenance phase.

In principle, PIP_2_ could alter PAR polarity during the maintenance phase through an impact on F-actin organization or else via an actin-independent role. A potential role of F-actin in polarity maintenance is somewhat controversial, in contrast to its well known role during polarity establishment (Goehring et al., 2011; Hill and Strome, 1990; Liu and Fletcher, 2006; Severson and Bowerman, 2003). In the light of our findings with PIP_2_ level alterations, we set out to directly test whether F-actin plays a role in polarity maintenance, first by adding Cytochalasin D to *perm-1(RNAi)* embryos expressing Lifeact::mKate-2 and GFP::PAR-2, after polarity establishment (Fig. 7A, B). Consistent with previous studies (Goehring et al., 2011; Hill and Strome, 1990), we found that Cytochalasin D addition at this stage does not alter the GFP::PAR-2 domain in a major manner (Fig. 7A, B; Movie 15). However, we found also that Cytochalasin D treatment does not fully disrupt F-actin, as clumps of Lifeact::mKate-2 remain present on the embryo anterior (Fig. 7B). We hence turned to inhibiting F-actin polymerization using Latrunculin A, which resulted in a total depletion of F-actin (Fig. 7C; N=12; Movie 16). We observed membrane invaginations that remove GFP::PAR-2 from the cortex into cytoplasmic aggregates (Fig. 7C, arrowhead), as reported by others (Goehring et al., 2011; Redemann et al., 2010). Importantly, in addition, we monitored changes in GFP::PAR-2 distribution not as a function of drug addition time, as in earlier work (Goehring et al., 2011), but of the time t_1/2_ at which half of the Lifeact::mKate-2 fluorescence disappeared from the membrane. In doing so, we found that the size of the GFP::PAR-2 domain decreased significantly after t_1/2_ in all embryos analyzed (Fig. 7C, bottom; N=12). As reported in Figure S5J, we found that the t_1/2_ of Lifeact::mKate-2 and of GFP::PAR-2 disappearance are highly correlated (Pearson coefficient p=0.86; p=0.0014). This finding reinforces the conclusion that F-actin is critical not only for the establishment, but also for the maintenance of PAR polarity.

**Fig. 7.**
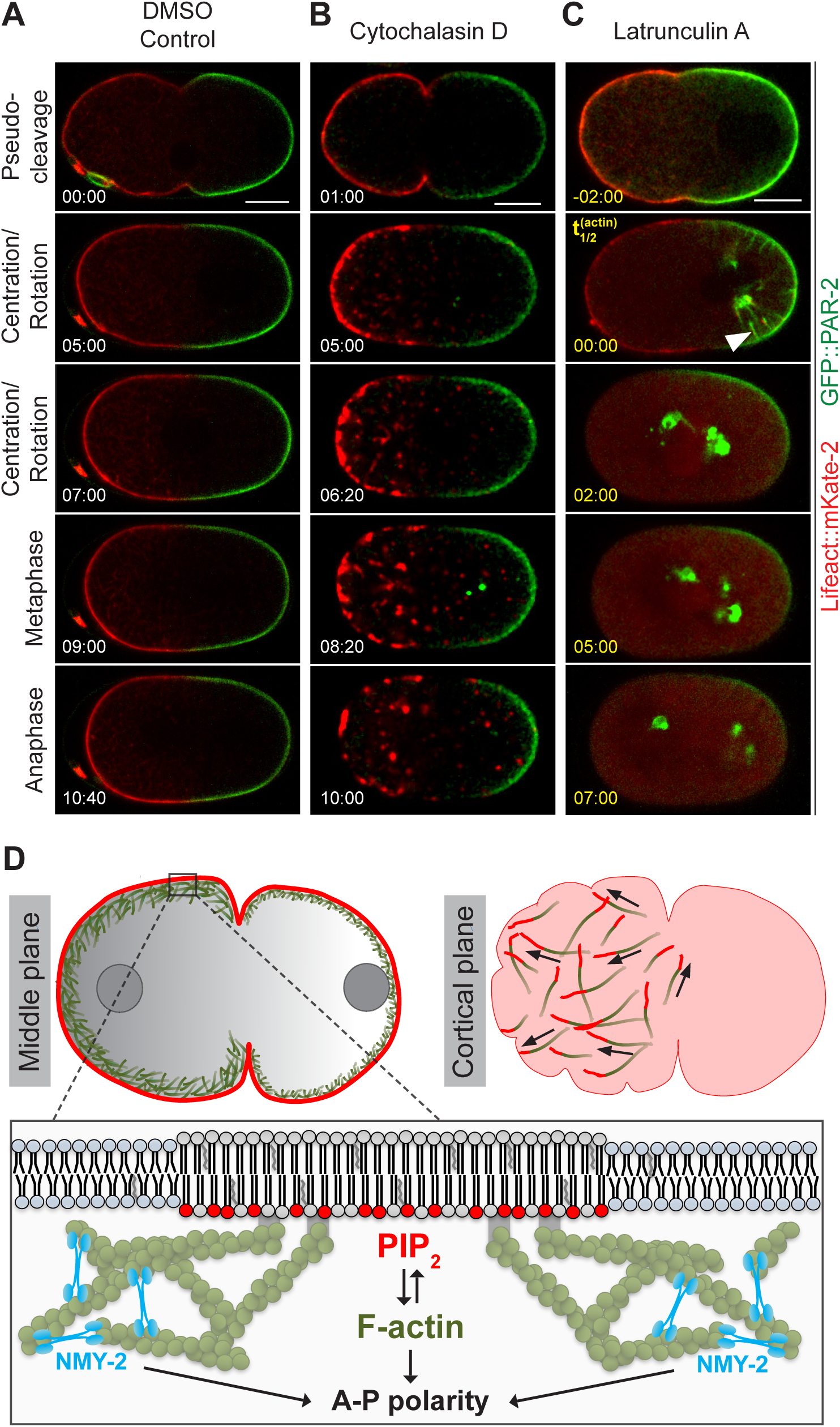
F-actin impairment affects GFP::PAR-2 also during polarity maintenance. **(A-C)** Confocai spinning disk cortical imaging of *perm-1(RNAi)* embryos expressing GFP::PAR-2and Lifeact::mKate-2 (middle plane), treated during early centration/rotation either with DMSO (A, N=6), Cytochalasin D (B, N=5, note that this movie was acquired with bining=2), or Latrunculin A (C, N=18). Arrowhead points to plasma membrane invagination (Redemann et al., 2010). See Movies 15, 16. **(D)** Graphical summary and working model (not to scale). PIP_2_ is enriched in dynamic and polarized structures at the cortex of one-cell *C. elegans* embryos. These structures move ahead of F-actin fibers, with which their velocity and direction is correlated. Moreover, the two components exhibit mutually reciprocal requirements, as the formation of PIP_2_ cortical structures requires F-actin, whereas a proper PIP_2_ level is essential for F-actin network organization. Moreover, through its ability to properly reorganize the F-actin network, PIP_2_ is essential for proper sizing of PAR domains and thus for A-P polarity establishment and maintenance. In addition, there might be an actin independent pathway through which PIP_2_ regulates polarity. See text for further details. Note that in the cortical embryo schematic on the top right, for simplicity only those F-actin filaments that move in concert with PIP_2_ cortical structures are represented.

Overall, we uncovered that a proper level of PIP_2_ is essential for correct sizing of PAR domains presumably through reorganization of F-actin, not only during polarity establishment but also during polarity maintenance phase.

## Discussion

In this work, we demonstrate that PIP_2_ forms cortical structures in the one-cell C. *elegans* embryo. We show that these structures depend on F-actin and, reciprocally, that PIP_2_ regulates F-actin organization, revealing an interdependence of these two components in the worm zygote (Fig. 7D). Moreover, likely through its impact on the actin cytoskeleton, PIP_2_ is also needed for the correct sizing of anterior and posterior PAR domains, demonstrating for the first time that a plasma membrane lipid component participates in setting A-P polarity in the *C. elegans* embryo.

### PIP_2_ is present in discrete cortical structures in C. elegans zygotes

The distribution and dynamics of PIP_2_ at the plasma membrane of early *C. elegans* embryos were not clear prior to this work, primarily because the middle embryo plane has been analyzed in most past investigations. Here, we developed cortical imaging conditions to assay subcellular distributions at the cortex, the very location where the function of PAR proteins and components critical for asymmetric division is exerted. In doing so, we discovered that PIP_2_ is present in dynamic polarized cortical structures marked by the PIP_2_ biomarker GFP::PH^PLC1δ1^, in line with recent observations mentioning a non-uniform distribution of this fusion protein (Rodriguez et al., 2017; Wang et al., 2017). Although patches of plasma membrane enriched in PIP_2_ have been observed in other systems (Chierico et al., 2015; Golub and Caroni, 2005; McLaughlin et al., 2002; Zhang et al., 2012), the stereotyped progression through the first cell cycle of the large *C. elegans* zygote enabled us to uncover their distribution and dynamics with unprecedented resolution. Why were such remarkable structures not observed in previous studies in the worm? In addition to the fact that they are not noticeable when imaging the middle plane of the embryo, other plausible reasons include that PIP_2_ cortical structures appear only transiently during the cell cycle and that they are not preserved upon fixation (data not shown).

How do PIP_2_ cortical structures assemble? We hypothesize that PIP_2_ cortical structures form by redistribution of existing PIP_2_ rather than by *de novo* synthesis through the PIP5K1 PPK-1, and this for two reasons. First, PIP_2_ in other systems has been suggested to diffuse much faster than it is synthesized (McLaughlin et al., 2002), such that potential local synthesis is unlikely to dictate restricted PIP_2_ localization. Second, PPK-1, the sole PIP5K1 in *C. elegans,* is enriched in the posterior of the embryo (Panbianco et al., 2008), away from the location where most PIP_2_ cortical structures are. Interestingly, we find also that PIP_2_ cortical structures form and move independently of actomyosin contractions, as evidenced by their unchanged presence and behavior upon NMY-2 depletion (data not shown). Nevertheless, it remains possible that PIP_2_ cortical structures form at membrane protrusions or ruffles, which would be consistent with the finding that PIP_2_ can stimulate F-actin polymerization in curved but not in flat membranes (Gallop et al., 2013), and with PIP_2_ accumulating in membrane ruffles, nascent phagosomes and the leading edge of motile cells (reviewed in McLaughlin et al., 2002; Zhang et al., 2012). Overall, we propose that, in the *C. elegans* zygote, PIP_2_ cortical structures form through the redistribution of existing PIP_2_ at the plasma membrane, perhaps preferentially at membrane protrusions or ruffles.

### Interdependence of PIP_2_ and F-actin

PIP_2_ and F-actin exhibit a reciprocal relationship in a number of systems, and we uncover here that this is the case also in *C. elegans* embryos. We found that PIP_2_ cortical structures and F-actin movements are coupled, with PIP_2_ structures moving slightly ahead of F-actin filaments, at velocities compatible with actin polymerization driving their movements. This leads us to propose that actin polymerization pushes PIP_2_ cortical structures, in a manner reminiscent of other actin-dependent motility processes such as that of *Listeria monocytogenes* (reviewed in Mogilner and Oster, 1996). While being pushed ahead of F-actin filaments in *C. elegans,* PIP_2_ structures might recruit factors promoting actin polymerization and branching, such as ECT-1, RHO-1 and CDC-42, thus guiding proper F-actin network reorganization, in line with suggestions in other systems (reviewed in Chichili and Rodgers, 2009). Intriguingly, the distribution of a biosensor that detects active RhoA overlaps with that of NMY-2 foci (Reymann et al., 2016; Tse et al., 2012). Given that we show here that PIP_2_ cortical structures do not overlap with NMY-2, while they do overlap with GFP::RHO-1, perhaps the bulk of RHO-1 associating with PIP_2_ cortical structures is not active. Alternatively, given that we show here that RHO-1 colocalizes with its activating GEF ECT-2, perhaps the RhoA biosensor used previously does not detected all active RHO-1 species. Furthermore, it is interesting to note that non-muscle myosin 2 plays a role in actin network disassembly in fish keratinocytes (Wilson et al., 2010). Extrapolating from this observation, it is tempting to speculate that PIP_2_, by promoting F-actin assembly, and NMY-2, by promoting F-actin disassembly, in addition to its role in network contractility, may together ensure proper F-actin dynamics in the early *C. elegans* embryo.

### PIP_2_ and PAR-dependent polarity

PAR proteins are also distributed unevenly within their cortical domain. For instance, PAR-6 exists in two cortical populations, one diffuse that depends on CDC-42, and one punctate that colocalizes with PAR-3 (Beers and Kemphues, 2006; Robin et al., 2014). Moreover, PAR-3 forms clusters that are crucial for proper polarity and that assemble in a manner dependent on PCK-3, CDC-42, as well as actomyosin contractility (Rodriguez et al., 2017; Wang et al., 2017). Intriguingly, we find that PIP_2_ cortical structures colocalize within the more diffuse cortical PAR-6 protein population, the one lacking PAR-3 (Beers and Kemphues, 2006; Robin et al., 2014; Rodriguez et al., 2017; Wang et al., 2017). We establish here that increasing the level of PIP_2_ augments the segregation of both PAR-6 populations to the embryo anterior, potentially because cortical clustering of PAR-3 depends on the actomyosin cytoskeleton and cortical tension (Beers and Kemphues, 2006; Robin et al., 2014; Rodriguez et al., 2017; Wang et al., 2017). Overall, we propose that increasing the level of PIP_2_ might augment cortical tension, which could in turn promote PAR-3 clustering and thereby aid segregation.

We show that a correct PIP_2_ level is essential for proper polarity establishment and maintenance trough correct positioning of GFP::PAR-2 and GFP::PAR-6 domains. When increasing the level of PIP_2_, the continued movement of PAR domains towards the anterior until the end of mitosis alters their relative size. This is reminiscent of changes in the size of PAR domains that occur upon RGA-3/4 depletion (Schonegg et al., 2007). However, although *rga-3/4(RNAi)* embryos exhibit a hypercontractile actomyosin network, this is not the case of embryos with increased PIP_2_ level. We thus propose that actomyosin contractility regulated by NMY-2 and F-actin organization regulated by PIP_2_ contribute in concert to correct movements of the actomyosin network, and, therefore, proper sizing of PAR-polarity domains.

### On the role of the actomyosin network in polarity establishment and maintenance

The actomyosin network plays a well-established role during polarity establishment, whereas its role during polarity maintenance has been less clear (Goehring et al., 2011; Hill and Strome, 1990; Liu and Fletcher, 2006; Severson and Bowerman, 2003). Our results, together with that of others, indicate that the actomyosin network regulates the size and localization of PAR domains in two ways. First, when the actomyosin network moves along the A-P embryonic axis, PAR domains alter their distribution accordingly. This relationship was clear prior to this work for the polarity establishment phase, and we show here that this is also the case during polarity maintenance. Second, actin has been suggested to play merely a passive role during the maintenance phase in preventing cortical PAR-2 removal through membrane invaginations driven by microtubules (Goehring *et al.,* 2011). We reveal here that the lack of this function can lead to the near total disappearance of cortical PAR-2, emphasizing the critical importance of actin also during the maintenance phase.

Overall, our results in the *C. elegans* zygote are consistent with the role of PIP_2_ in F-actin reorganization and polarity in other organisms. Previous work in C. *elegans* showed that depletion of CSNK-1, which negatively regulates PPK-1 localization, does not influence polarity at the end of the first cell cycle (Panbianco et al., 2008). Perhaps PPK-1 distribution plays only a minor role in regulating the cellular level of PIP_2_ in the zygote, being dispensable for PIP_2_ cortical structure formation. In this case, depleting an enzyme such as CSNK-1 that negatively regulates PPK-1 localization would not be expected to influence PIP_2_ cortical distribution. Here, by contrast, we establish unequivocally that alterations in the level of PIP_2_ impairs polarity establishment and maintenance during the first asymmetric division of *C. elegans* embryos.

## Material and Methods

### Worm Strains

Nematodes were maintained at 24°C using standard protocols (Brenner, 1974). The following worm strains were used: GFP::PH^PLC1δ1^ (OD58, *unc-119(ed3)* III; ltIs38[*pie*-*1*p::GFP::PH(PLC1delta1) + *unc-119(+)])* (Audhya et al., 2005); mCherry::PH^PLC1δ1^ (OD70, *unc-119(ed3)* III; ltIs44[p/e-1p::mCherry::PH(PLC1delta1) + *unc-119(+)]V)* (Audhya et al., 2005); GFP::PAR-2 (TH129) and GFP::PAR-6 (TH110) (Schonegg et al., 2007); GFP::NMY-2 (LP162, *nmy-2(cp13[nmy-2::gfp* + LoxP]) I) (Dickinson et al., 2013); CAV-1::GFP (RT688, unc-119(ed3) III; pwIs281[CAV-1::GFP, *unc-119(+)])* (Sato et al., 2006); mNeonGreen::PH^PLC1δ1^ (LP274, cpIs45[P*mex*-5::mNeonGreen::PLCδ-PH::*tbb*-2 3’UTR + *unc-119(+)]* II; *unc-119(ed3)* III), mKate-2::PH^PLC1δ1^ (LP307, cpIs54[P*mex*-5::mKate2::PLCδ-PH(A735T)::*tbb*-2 3’UTR + *unc-119*(*+*)] II; *unc-11*9(ed3) III) and mCherry::PH^PLC1δ1^ (LP308, cpIs55[P*mex*-5::mCherry-C1::PLCδ-PH::*tbb*-2 3’UTR + *unc-119*(*+*)] II; *unc-119*(ed3) III) (Heppert et al., 2016); Lifeact::mKate-2 (strain SWG001) (Reymann et al., 2016); GFP::RHO-1 (SA115, *unc-119(ed3)* III; tjls1[*pie-1*::GFP::*rho*-*1* + *unc-119(+)])* (Motegi et al., 2006); GFP::CDC-42 (SA131, *unc-119(ed3)* III; tjIs6*[pie-1p::GFP::cdc-42 + unc-119(+)].)* (Motegi and Sugimoto, 2006); GFP::ECT-2 (SA125, unc-119(ed3) III; tjls4[*pie*-*1::*GFP*::ect-2+unc-119(+)])* (Motegi and Sugimoto, 2006); *unc-26(s1710)* (EG3027, *unc-26(s1710)* IV) (Charest et al., 1990); *age-1(m333),* (DR722, age-1(m333)/mnC1 *dpy-10(e128) unc-52(e444)* II) (Riddle, 1988). Crosses of worm strains were performed at 20°C to generate lines homozygote for all transgenes, which were then maintained at 24°C. For GFP::RHO-1 and mCherry::PH ^PLC1δ1^, as well as GFP::PH^PLC1δ1^ and Lifeact::mKate-2, worm lines were crossed and F1 progeny heterozygote for both transgenes imaged.

### RNAi

RNAi-mediated deletion was performed essentially as described (Kamath et al., 2001), using bacterial feeding strains either from the Ahringer (Kamath et al., 2003) or the Vidal library (Rual et al., 2004) (gift from Jean-François Rual and Marc Vidal). RNAi for *par-2* (Ahringer), *par-3* (Ahringer), *nmy-2* (Ahringer), *act-1* (Vidal), *tba-2* (Vidal), and *ocrl-1* (Ahringer) was performed by feeding L3-L4 animals with bacteria expressing the corresponding dsRNA at 24°C for 20-26 hours. RNAi for *perm-1* (Ahringer) was performed by feeding L4 and young adults with bacteria expressing *perm-1* dsRNA at 20°C for 12-18 hours. The effectiveness of the deletion was screened phenotypically as follows: *par-2(RNAi)* and *par-3(RNAi)* -symmetric spindle positioning and cell division; *nmy-2(RNAi)* and *act-1(RNAi)* -absence of cortical ruffles, symmetric spindle positioning, no cytokinesis; *tba-2(RNAi)* -defective pronuclear meeting, no centration/rotation, no spindle assembly, misplaced cytokinesis furrow specification; *ocrl-1(RNAi)* -see results, *perm-1(RNAi):* successful action of added drug.

### Live imaging

Gravid hermaphrodites were dissected in osmotically balanced blastomere culture medium (Shelton and Bowerman, 1996) and the resulting embryos mounted on a 2% agarose pad. DIC time lapse microscopy (Fig.1 A,C,E,G,I) was performed at 25°C ± 1 °C with a 100× (NA 1.25 Achrostigmat) objective and standard DIC optics on a Zeiss Axioskop 2 microscope. All other images were acquired using an inverted Olympus IX 81 microscope equipped with a Yokogawa spinning disk CSU - W1 with a 63 (NA 1.42 U PLAN S APO) objective and a 16-bit PCO Edge sCMOS camera at 23°C. Images were obtained using 488 nm and 561 nm solid-state lasers with an exposure time of 400 ms and a laser power of 20-60%. For cortical imaging, 3 planes at the cell cortex (each 0.25 µm apart) were acquired. Cell cycle stages were determined using transmission light microscopy by imaging the middle plane in parallel (data not shown).

### Image processing and analysis

Cortical images of GFP::PH^PLC1δ1^ used for quantification were processed as follows: the 3 cortical planes were z-projected using average intensity projection, then a median filter of 1 pixel was applied. The background of the entire image was subtracted using the measured mean background in each frame. Signal intensity decay due to photobleaching was corrected with the Fiji plugin "bleach correction" using the exponential fitting method. The entire cortical region was segmented by applying a binary automated histogram-based threshold, followed by iterated morphological operations. Cortical structures were segmented by applying a binary intensity threshold, calculated by fitting the pixel intensity histogram with a Gaussian function and setting the threshold at 4 sigma from the Gaussian peak. The size of cortical structures was normalized to the total cortical area.

Curves of normalized cortical structures sizes were fit with a sigmoidal model and synchronized, setting the sigmoid inflection point, which corresponded typically to the time of centration/rotation, as time t=0 s. Curves of normalized cortical structures sizes were aligned manually for *act-1(RNAi)* and *unc-26(s1710) ocrl-1(RNAi)* embryos using the clear landmark provided by Nuclear Envelope Breakdown (NEDB) as a reference, because a sigmoid function could not be fit with the PH markers in these cases. Since the time separating centration/rotation from NEDB is typically150 seconds, t=0 was set at −150 seconds prior to NEDB for *act-1(RNAi)* and *unc-26(s1710) ocrl-1(RNAi)* embryos.

The Elongation Index was calculated as follows: ((perimeter^˄^2)/area)/4π using the MATLAB image processing function "regionprops". We then normalized the Elongation Index by a factor of 1/π such that a square of 2 × 2 pixels has an Elongation Index of 1.

Cortical images obtained by live confocal spinning disk imaging and shown in the figures were processed as follows: the 3 cortical planes were z-projected using maximum intensity projection, then a median filter of 1 was applied. The grey value fluorescence intensity of some transgenes (GFP::PH^PLC1δ1^ as heterozygote, GFP::PAR-2, GFP::PAR-6, Lifeact::mKate-2, mNeonGreen-PH^PLC1δ1^) was slightly variable likely resulting from variable expression/folding of the fluorescent fusion protein, which may stem for several reasons, including F1 heterozygosity, temperature shifting, silencing during crossing. The brightness and contrast of images resulting from embryos expressing these transgenes was therefore adjusted accordingly. The fluorescence intensity of mCherry::PH^PLC1δ1^ was also variable in some cases; therefore, the brightness and contrast of images from embryos expressing mCherry::PH^PLC1δ1^ were therefore adjusted accordingly as well. Such variability was especially pronounced in UNC-26 and OCRL-1 depleted embryos. To compare the intensity of mCherry-PH^PLC1δ1^ in control embryos and *unc-26(s1710) ocrl-1(RNAi)* embryos, 3 cortical planes, acquired as described above, were z-projected by summing the intensity of all slices. The resulting mean intensities were then computed as follows. First, a Otsu threshold was used to retrieve the brightest elements - including the embryo - of the image, retaining only the biggest blob, corresponding to the embryo. Values outside the embryo were averaged to obtain the mean background intensity value, which was subtracted from the embryo pixels. Thereafter, embryo pixel values were averaged to obtain the mean pixel intensity value.

### Cortical flow measurement, correlation analysis, and PIP_2_ structures tracking

For Particle Image Velocimetry (PIV) analysis, cortical image sequences of mNeonGreen::PH^PLC1δ1^ and Lifeact::mKate-2 were prepared by performing a maximum intensity z-projection of a stack of 2 planes (0.25 µm apart) and applying a median filer of 1 pixel. PIP_2_ cortical structures and the F-actin network were then segmented using the following procedure: the embryo was first extracted from the background using a histogram-based automated threshold, keeping only blobs of a size superior to one third of the biggest blob. The resulting binary images were deemed to be the embryo area. We applied a morphological erosion to the mNeonGreen::PH^PLC1δ1^ movies with a large structuring element (a disk 30 pixels in radius) to calculate the average value of the pixels not corresponding to PIP_2_ cortical structures; the PIP_2_ cortical structures were then segmented as the pixels of intensity higher than the computed average value, times a scaling factor determined empirically (1.7). The extraction of the F-actin network was achieved simply by determining a histogram-based automated threshold on the morphological top-hat of the F-actin image. F-actin filaments and PIP_2_ cortical structures where segmented prior to PIV analysis to ensure that only flow fields in the region of interest are measured.

PIV was then performed to measure cortical flows using the MATLAB based PIVlab toolbox (Thielicke and Stamhuis, 2014); this splits each image of a movie into a regular grid, for which the size of grid cells is given by the user. The position of each cell in the next image is estimated by finding the maximum normalized cell-to-cell cross-correlation of equivalent sizes in a geometrical neighborhood called interrogation area. PIV was applied to mNeonGreen::PH^PLC1δ1^ and Lifeact::mKate-2 separately, after segmentation of the corresponding cortical structures. The choice of the sizes of the cells and interrogation areas was a balance between two criteria: smaller cells allow to compute displacements with high spatial resolution, but excessively small cells do not contain enough information to be reliably correlated to other cells; the estimation of displacements of bigger cells is hence more reliable, but are computed with lesser resolution. We found empirically that 32 × 32 pixels for cell sizes, and 64 × 64 pixels for interrogation areas, to be a good compromise.

The PIV velocity fields output for both mNeonGreen::PH^PLC1δ1^ and Lifeact::mKate-2 signals were compared in terms of angles between colocalized features and correlation of the norms. For each movie, angles between velocity vectors of colocalized features were computed and plotted on a histogram. The average angle value for each time point and each movie was also computed, so as to monitor the coherence between the two vector fields over time. Similarly, we computed the correlations of the norms of all velocities in the two movies, for the whole movies, and also time-wise. The cut-off angle is defined as the θ 0 parameter of the curve of equation y = a exp(-θ/θ 0) fitted to the histogram.

Cross-correlation analysis was performed as follows. Movies used to calculate the cross-correlation were acquired alternating acquisition of red or green channel first to prevent introducing a bias through the order of image acquisition. The colocalization of the thresholded PIP_2_ cortical structures and F-actin network for a variety of time shifts was computed considering a time shift Δt (positive or negative), the colocalization of the segmented PIP_2_ image at time t and the segmented F-actin image at time t-Δt using the following formula:

Colocalization = (PIP_2_(t) ∩ F-actin(t-Δt)/ PIP_2_ (t)

Colocalization was computed in this manner from Δt =-(T-1) to Δt = (T-1), where T is the total duration of the movie. The Δt for which Colocalization is maximal represents the time shift between PIP_2_and F-actin. The mean time shift and its error were computed as follows: we fitted a parabola of equation y = a + (t-t0)^˄^2 + b to the location correlation as a function of the time shift. We calculated the best a, b and t0 parameters using a least-squares method, and input the standard deviations of the correlations to create a weight matrix used during the adjustment. The results were the mean time shift t0 = 9.3s and the standard deviation sigma_t0 = 1.5s.

To track PIP_2_ structures, embryos expressing mNG-PH^PLC1δ1^ were imaged with an exposure time of 50 ms, laser power of 60% and 70 ms frame rate. PIP_2_ structures were tracked manually on maximum intensity z-projection of the images containing the moving PIP_2_ structures of interest. The length of the track was obtained by reslicing it using the Fiji plugin “Reslice”. Velocity was calculated from the corresponding number of time points and track length.

### Drug addition

The eggshell was permeabilized by performing *perm-1(RNAi)* as described above. Gravid hermaphrodites were dissected in a cell culture dish with a glass bottom, and the resulting embryos imaged with an inverted confocal spinning disk microscope (see above). Drugs were added under the microscope while imaging to precisely control the timing of drug addition. The following drugs and concentrations were utilized: 30 µM Ionomycin (Calbiochem, 407950), 3-5 mM CaCl_2_ (Sigma-Aldrich, C5080), 20 µM Cytochalasin D (AppliChem, 22144-77-0), 12.5 µM Latrunculin A (Sigma-Aldrich, 76343-93-6). For control movies, DMSO at a concentration equivalent to the final DMSO concentration in the drug solutions was added to the buffer prior to dissection.

Successful drug action was determined for each embryo by the disappearance of the PH^PLC1δ1^ fluorescence signal from the plasma membrane (Ionomycin/ Ca^2+^, Latrunculin A) and of Lifeact::mKate-2 from the cell cortex (Latrunculin A, Cytochalasin D). The time between drug addition and drug action was variable, probably due to variations in eggshell permeability upon *perm-1(RNAi).* As a comparable reference time between embryos, we therefore determined the time t_1/2_ (t inflection) when half of fluorescence at the plasma membrane has disappeared. t_1/2_ was determined as follows: the total cortical region of the embryo was segmented by applying a binary automated histogram-based threshold. Fluorescent values at a distance of 20 pixels from the edge were measured, and their mean fluorescence values plotted over time; the inflection point of a fitted sigmoid function was then determined as t_1/2_.

### Lipid delivery

BODIPY^®^ FL Phosphatidylinositol 4,5-bisphosphate (Echelon Bioscience, C-45F6) (end concentration: 2 µM) was delivered to *perm-1(RNAi)* embryos by adding it to the buffer in which gravid worms were dissected. It typically helped to premix BODIPY^®^ FL Phosphatidylinositol 4,5-bisphosphate (100 µM) with a carrier histone (P-9C2; Echelon Bioscience) (300 µM) (Ozaki et al., 2000). Phosphoinositide-histone complexes were formed as described (Kotak et al., 2014): both components were mixed by vigorous vortexing and then incubated for at 15 min at room temperature.

### Statistical analyses

The software package JMP 13.2.0 (SAS Institute GmbH) and MATLAB 2016 were used to perform statistical analysis. Two-group comparisons were performed using Student’s t-test. Results with values of p≤0.05 were considered statistically significant.

## Acknowledgments

We thank Kalyani Thyagarajan and Sachin Kotak for initial observations of PIP_2_ cortical structures, Olivier Burri (Biolmaging and Optics Platform, BIOP, School of Life Sciences, EPFL) for help in developing the script for preprocessing of embryos with ImageJ, as well as the BIOP at large for microscopy support. For strains, we thank John Audyha, Daniel Dickinson, Bob Goldstein, Barth Grant, Stephan Grill, Anthony Hyman, Karen Oegema, and Anne-Cécile Reymann, as well as the *Caenorhabditis* Genetics Center (CGC), which is funded by NIH Office of Research Infrastructure Programs (P40 OD010440). This work was supported by the Swiss National Science Foundation (31003A_155942). The funders had no role in study design, data collection and analysis, decision to publish, or preparation of the manuscript.

